# Cross-Recording Handwritten Digit Decoding from sEMG Using a Compact CNN–Transformer and Few-Shot Adaptation

**DOI:** 10.64898/2026.08.12.740174

**Authors:** Anna V. Makarova, Maria V. Golitsyna, Mikhail A. Lebedev

## Abstract

Surface electromyography (sEMG) offers a silent and wearable input modality, but its practical use is limited by variability across users and recording sessions. This study presents a compact CNN– Transformer model for decoding isolated handwritten digits from eight-channel sEMG signals. The model combines trainable signal preprocessing, convolutional feature extraction, and Transformer-based temporal modeling. It was evaluated on ten recordings from five participants using recording-seen classification, leave-one-recording-out (LORO) generalization, and few-shot adaptation. The model achieved a mean macro F1 score of 0.924 *±* 0.059 in the recording-seen setting and 0.619 *±* 0.252 under zero-shot LORO evaluation. Adaptation using two labeled trials per digit increased macro F1 to 0.828 *±* 0.112, while ten trials per digit achieved 0.925 *±* 0.053. The proposed architecture also outperformed classical and neural baselines in the controlled LORO benchmark. These results indicate that compact CNN–Transformer models, combined with lightweight target-recording calibration, provide a promising basis for adaptive sEMG-based input systems.

## 1 Introduction

Surface electromyography (sEMG) is a non-invasive method to record muscle activity, which has become an important signal modality in human–computer interaction, assistive technologies, prosthetic control, and recognition of human activity [1]. In contrast to conventional handwriting recognition, where the model usually receives a visual image of a written character trajectory, EMG-based handwriting decoding infers written symbols directly from neuromuscular signals. This makes sEMG handwriting decoding relevant as a potential input component in multimodal interfaces. It is also relevant for silent input systems and for assistive interfaces where mechanical keyboards, touchscreens, or visually guided hand tracking are inconvenient, unavailable, or physically demanding.

In this work, handwritten input is understood as the use of writing movements to compose symbolic input through an interface, while EMG-based handwritten input refers specifically to translating EMG signals recorded during writing into textual symbols. This formulation is important because the input is not the written trace itself, but the muscle activity that precedes and accompanies handwriting. As a result, the recognition problem differs from image-based optical character recognition: the model must learn temporal neuromuscular patterns associated with intended handwriting patterns rather than visual shapes.

Prior work has shown that handwriting-related EMG signals can be decoded using machine learning and deep learning methods. A central contribution in this area is the HCMYO-A line of work, where Beltran-Hernandez et al. proposed multi-stroke handwriting character recognition from sEMG using convolutional-recurrent neural networks [2]. Their work demonstrated that sEMG contains discriminative information for recognizing handwritten digits and Latin letters and that neural architectures can learn useful temporal representations from such signals. The associated HCMYO-A dataset includes sEMG recordings for 36 character classes, including digits and Latin letters, collected during handwriting activity [3]. This dataset is especially relevant because it frames handwriting recognition from EMG as a sequence-recognition task rather than a static classification problem.

The sequential nature of handwriting is one of the main reasons why EMG-based decoding is challenging. A written digit or character is not produced by a single instantaneous muscle activation pattern, but by a temporally extended sequence of coordinated contractions. The same symbol may be written with different stroke order, speed, pressure, and motor strategy across users. Even within the same user, writing style may vary across repetitions because of fatigue, attention, posture, or electrode contact. Therefore, an effective model must capture both local signal patterns and longer temporal dependencies. Earlier convolutional-recurrent approaches address this by combining convolutional layers for feature extraction with recurrent units such as LSTM or GRU for sequence modeling [2]. This design is well-suited for handwriting because local EMG fluctuations can encode short motor primitives, while recurrent layers can represent their temporal organization.

However, recurrent architectures are not the only way to model temporal structure. The Transformer architecture employs self-attention as the mechanism for modeling dependencies across sequence positions without relying on recurrence [4]. sEMG gesture-recognition studies have demonstrated the utility of Transformer-based temporal modeling [5, 6], motivating its evaluation for handwriting-related signals. A CNN-Transformer architecture can therefore combine two complementary inductive biases: convolutional layers can extract local temporal and channel-wise features from raw or minimally processed EMG while Transformer blocks can model relationships between distant time steps and dynamically weight the most informative parts of the sequence. This hybrid design is particularly suitable for handwriting recognition, where both short local activations and global stroke-level structure matter.

Signal preprocessing remains an important part of EMG-based recognition pipelines. sEMG signals are noisy, non-stationary, and sensitive to acquisition conditions, so filtering and normalization are commonly used before model training. Classical Butterworth filters are frequently used to suppress unwanted frequency components while preserving the shape of the relevant signal band [7]. In deep learning pipelines, preprocessing should be strong enough to reduce artifacts but not so aggressive that it removes discriminative temporal information needed by the model.

A major limitation of EMG-based handwriting recognition is generalization across users and recording sessions. sEMG patterns depend on individual anatomy, skin impedance, electrode placement, muscle fatigue, writing posture, and individual motor strategy. Consequently, models trained on one group of users’ recordings often show degraded performance when applied to unseen subjects. This issue is well documented in the broader field of myoelectric pattern recognition. Zhang et al. studied multi-source domain generalization and adaptation for cross-subject myoelectric recognition and emphasized that individual differences can produce substantial distribution shifts between training and target users [8]. This problem is directly relevant to EMG handwriting decoding: a model that performs well under within-recording or randomly split evaluation may still fail in a realistic scenario where a new user must be recognized with little or no calibration data.

For improving decoder generalization, evaluation protocols are as important as model architecture. Leave-one-recording-out evaluation, which includes both cross-participant and within-participant cross-session transfer, provides a stricter test of whether the model learns handwriting representations transferable across recordings. In addition, few-shot or low-data adaptation protocols are practically important because fully calibrating a model for every new user could be time-consuming. Fine-tuning on a small fraction of data from a new user can be viewed as a compromise between generic and personalized modeling. Related few-shot learning methods, such as prototypical networks, show how models can be designed to generalize from limited examples by learning an embedding space where class prototypes are informative [9].

Recent progress in deep learning has shifted EMG recognition from handcrafted features toward end-to-end or representation-learning approaches [1]. This trend is promising because raw EMG contains complex temporal, spectral, and spatial information that could be difficult to summarize using manually designed features alone. CNN-based layers can learn local filters directly from the signal, while attention-based mechanisms can learn which temporal segments are most relevant for classification. For handwritten digits, where class differences may be subtle and temporally localized, this is especially useful.

The present study evaluated a compact CNN–Transformer for isolated handwritten-digit decoding from sEMG under three complementary conditions: recording-seen classification, leave-one-recording-out generalization, and supervised adaptation using a limited number of labeled trials from the target recording. The study compared this architecture with the classical and neural baselines and examined its key components through controlled ablations. The main contributions of this study are fourfold: (1) introduction of a compact CNN–Transformer architecture for isolated handwritten-digit decoding from eight-channel sEMG; (2) a cross-recording evaluation that explicitly includes both cross-participant and within-participant cross-session transfer; (3) a systematic assessment of supervised target-recording adaptation under low-data calibration conditions; and (4) a controlled comparison with classical and neural baselines accompanied by component-level ablation analysis.

## 2 Methods

### 2.1 Dataset

We conducted handwritten character decoding using an existing sEMG dataset [10]. The analysis assessed ten recordings obtained from five participants. Four participants contributed one recording each, identified as A, B, C, and J, whereas participant M contributed six recordings on different days, identified as M1–M6. Each recording contained eight channels of sEMG acquired from forearm and hand muscles at a sampling frequency of 1 kHz while the participant repeatedly wrote the digits 0–9 on a graphics tablet. Each digit was written approximately 50 times per recording. The dataset also contained synced tablet coordinates, an indicator of contact between the pen and the writing surface, digit labels, and trial identifiers. These auxiliary signals were used to identify individual handwriting trials and visualize the written trajectories. Only the eight-channel sEMG signals were used as input to the classification models. The electrode locations and representative digit trajectories are shown in Figure 1.

**Figure 1:**
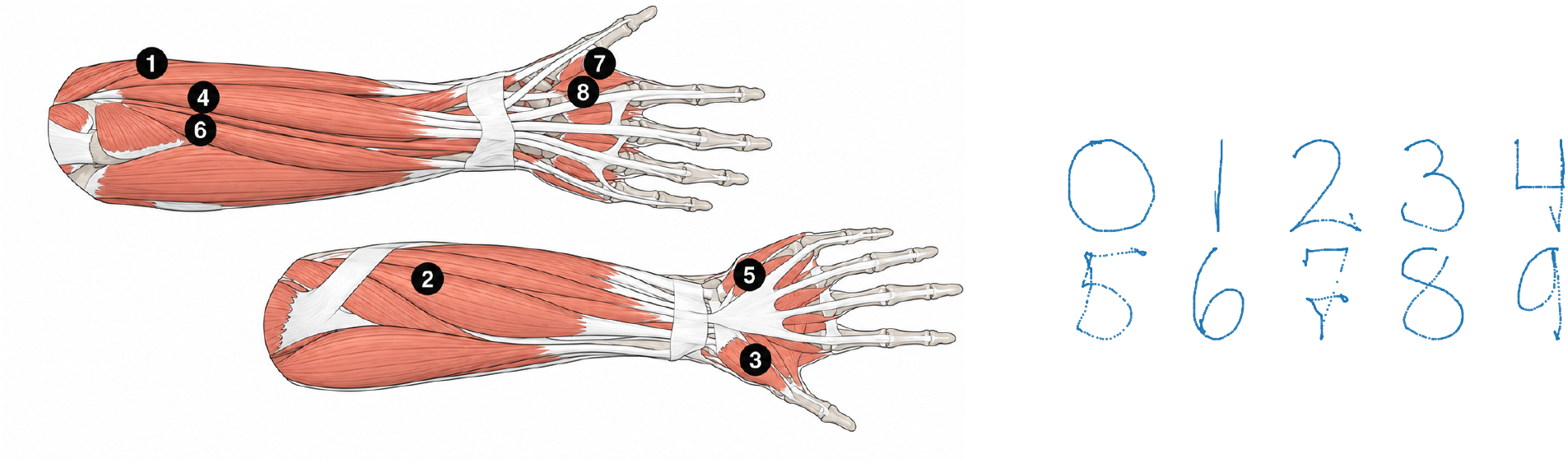
Locations of the 8 sEMG electrodes on forearm and hand muscles (ECR, FCR, OP, ED, APB, ECU, lFDI, mFDI) (left), and examples of handwritten digits (0–9) produced by participants during recording (right).

### 2.2 Signal Preprocessing

Continuous EMG recordings were filtered using a fourth-order zero-phase Butterworth band-pass filter (1–450 Hz), followed by a 50 Hz notch filter (*Q* = 30) to suppress power-line interference. Individual handwriting trials were extracted using the trial identifiers provided in the dataset. The active writing interval was defined by the *OnPaper* signal, indicating pen contact with the tablet surface. Trials without active handwriting or with extreme durations (above the 99th percentile) were discarded. To preserve the movement onset and offset, a 100 ms margin was added before and after each active interval. Each trial was resampled to a fixed length of 2500 samples using Fourier-domain resampling, producing a fixed-length representation that reduces variation in trial duration. The corresponding tablet trajectories were resampled using the same procedure to preserve temporal alignment. After preprocessing, each EMG sample was represented as an 8 *×* 2500 tensor, where eight is the number of EMG channels. The digit label for each trial was determined from the corresponding annotation in the original dataset. Tablet coordinates and auxiliary signals were used only for trial segmentation and visualization; all classification experiments used only the eight-channel sEMG signals as model input. Normalization was performed independently within each experimental split. Signal amplitudes were first clipped to the range [−6, 6]. For each EMG channel, the median and robust scale, defined as 1.4826 times the median absolute deviation, were estimated exclusively from the training subset across trials and time samples.

The resulting parameters were subsequently applied unchanged to the validation, support, and held-out test subsets. Therefore, no statistics from the held-out recording or its evaluation trials were used for normalization.

### 2.3 Proposed CNN–Transformer Model

A compact CNN–Transformer model was developed to classify handwritten digits directly from multichannel sEMG signals (Figure 2). The network combines trainable EMG preprocessing, convolutional feature extraction, and Transformerbased temporal modeling. The primary model used throughout this study employed a single-scale preprocessing block, while an alternative multiscale preprocessing variant was evaluated separately in the baseline and ablation experiments. Unless stated otherwise, the following description refers to the single-scale model.

**Figure 2:**
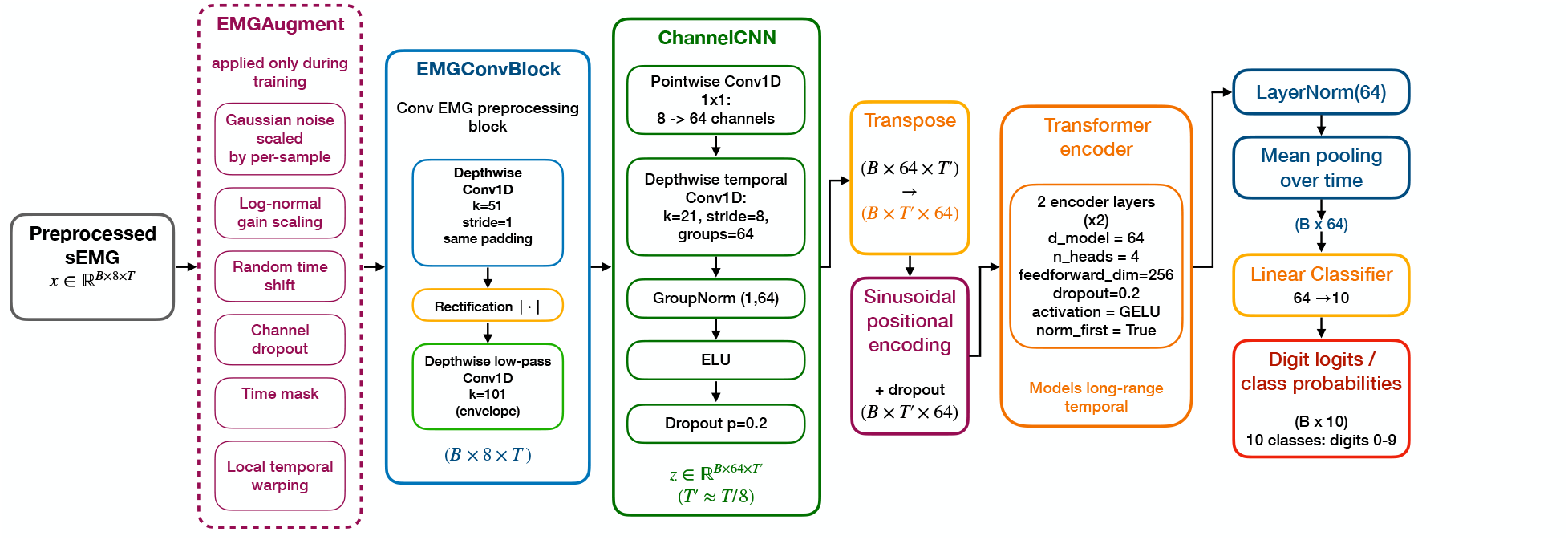
Architecture of the proposed single-scale CNN–Transformer for isolated handwritten-digit classification from preprocessed sEMG. A trainable depthwise filtering–rectification–envelope block is followed by pointwise channel mixing and strided temporal convolution. Sinusoidal positional encoding is added to the downsampled sequence before two Transformer encoder layers. Temporal mean pooling and a linear classifier produce logits for the ten digit classes.

#### 2.3.1 Trainable EMG Preprocessing

The input to the network was an eight-channel EMG segment of size 8 *×* 2500. The first processing stage consisted of a trainable preprocessing block designed to learn channel-specific temporal filtering followed by rectification and envelope-like smoothing. Each channel was independently processed by a depthwise one-dimensional convolution with a kernel size of 51, followed by absolute-value rectification and a second depthwise convolution with a kernel size of 101 that produced a learnable signal envelope.

#### 2.3.2 Local feature extraction

The preprocessed signals were passed to a convolutional feature extractor. A pointwise (1 *×* 1) convolution projected the eight input channels into a 64-dimensional feature space by learning linear combinations of muscle activations. A depthwise temporal convolution with a kernel size of 21 and stride 8 then extracted local temporal patterns while reducing the sequence length from 2500 samples to approximately 313 tokens. Group normalization, ELU activation, and dropout with a probability of *p* = 0.2 were applied after the depthwise temporal convolution.

#### 2.3.3 Transformer encoder and classifier

The resulting feature sequence was transposed to the (*B, T*, 64) representation and augmented with sinusoidal positional encoding. Long-range temporal dependencies were modeled using a two-layer Transformer encoder with an embedding dimension of 64, four attention heads, a feed-forward dimension of 256, GELU activation, pre-layer normalization, and dropout of 0.2. The encoded sequence was averaged over the temporal dimension and passed through layer normalization and a fully connected layer that produced logits for the ten handwritten digit classes.

#### 2.3.4 Multiscale Preprocessing

To investigate whether multiple temporal receptive fields improve recognition, an alternative preprocessing block was evaluated. In this variant, the initial depthwise convolution was replaced by three parallel depthwise convolutions with kernel sizes of 21, 51, and 71 samples. Their outputs were averaged before rectification and envelope extraction, while all subsequent network components and training procedures remained identical.

**Table 1:** Architecture summary.

| Component | Single-scale | Multiscale |
| --- | --- | --- |
| EMG Preprocessing | 1,216 | 1,952 |
| Local CNN | 1,984 | 1,984 |
| Transformer Encoder | 99,968 | 99,968 |
| LayerNorm | 128 | 128 |
| Classifier | 650 | 650 |
| **Total Parameters** | **103,946** | **104,682** |

**Table 2:** Augmentation parameters.

| Augmentation | Probability | Magnitude |
| --- | --- | --- |
| Additive Gaussian noise | 0.35 | SD = $0.10 \times$ sample RMS |
| Gain perturbation | 0.70 | $\exp(\mathcal{N}(0, 0.25^2))$ |
| Temporal shift | 0.50 | Up to 120 samples |
| Channel dropout | 0.45 | Channel-drop rate 0.25 (two of eight channels) |
| Temporal masking | 0.60 | 100 samples |
| Local temporal warping | 0.005 | Strength 0.30 |

#### 2.3.5 Model Training

All neural models were trained using the AdamW optimizer with an initial learning rate of 0.001 and a weight decay of 1 *×* 10−4 [11]. Learning-rate scheduling was performed using cosine annealing with *T*max = 50. Gradient clipping with a maximum norm of 1.0 was applied throughout training. Mini-batches contained 64 trials. During few-shot adaptation, the learning rate was reduced to 1 *×* 10−4, while all model parameters remained trainable. Model-specific training schedules and validation strategies are described in the corresponding experimental protocols.

#### 2.3.6 Data Augmentation

To improve robustness and reduce overfitting, stochastic data augmentation was applied during training. Each augmentation was enabled independently with a predefined probability, while validation and test samples were left unchanged. The augmentation pipeline included additive Gaussian noise, multiplicative gain perturbation, temporal shifts, channel dropout, temporal masking, and local temporal warping. These transformations were designed to simulate realistic variability in EMG acquisition, including sensor noise, changes in signal amplitude, temporal misalignment, and differences in handwriting dynamics, while preserving the underlying motor pattern.

### 2.4 Experimental Protocols

Three complementary evaluation protocols were designed to assess within-recording classification, cross-recording generalization, and rapid adaptation to unseen recordings.

#### 2.4.1 Within-Recording Evaluation

The within-recording protocol was used to estimate recording-seen classification performance when training and testing data originated from the same recording session. In each split, a single model was trained on the pooled training subsets from all recordings and evaluated on the pooled held-out subsets from the same recordings. Five independent stratified 90/10 train–test splits were generated for each recording using random seeds. To examine the influence of model selection, two training configurations were evaluated: fixed training for 60 epochs with no validation and training for a maximum of 60 epochs with 10% of the training data reserved for validation. In the latter case, the model checkpoint with the highest validation macro F1 score was selected for evaluation.

#### 2.4.2 Leave-One-Recording-Out Evaluation

To evaluate cross-recording generalization, we adopted a leave-one-recording-out (LORO) protocol. In each fold, one recording was excluded from training and used exclusively for testing, while all remaining recordings formed the training set. No samples from the held-out recording were used during model training or model selection. To investigate the influence of training duration on zero-shot generalization, models were trained for 30, 45, or 60 epochs without validation. Because participant M contributed multiple recordings acquired on different days, the LORO protocol evaluated both transfer across recording sessions of the same participant and transfer between different participants. As a control analysis, we additionally repeated training using sampling weights balanced across the five participants rather than across individual recordings to reduce the influence of repeated recordings.

#### 2.4.3 Few-Shot Adaptation

To evaluate rapid personalization for a previously unseen recording, the model was first pretrained on all non-target recordings for 30 epochs. A small support set sampled from the target recording was then used for supervised fine-tuning, while all remaining trials from the target recording were reserved exclusively for evaluation. Support sets containing 1, 2, 5, or 10 trials per digit were investigated. For one-shot and two-shot adaptation, fine-tuning durations ranging from 20 to 500 epochs were evaluated to analyze convergence behavior. Five-shot and ten-shot adaptations were evaluated using short (20 epochs) and extended (500 epochs) fine-tuning schedules.

#### 2.4.4 Validation Strategies

Selecting the optimal model for an unseen recording is challenging because validation data from the target recording are typically unavailable. We therefore evaluated two validation strategies that used only non-target recordings. In both strategies, the checkpoint with the highest validation macro-averaged F1 score was selected for testing, and early stopping with a patience of 15 epochs was applied. In the first strategy, one recording was held out exclusively for testing while the trials from each remaining recording were independently divided into training and validation subsets using a stratified split. Ten percent of the trials from each training recording were assigned to the validation set, and the remaining trials were used for model training. The model was trained for up to 55 epochs. This strategy evaluated whether model selection could be performed using validation trials drawn from the same recordings that contributed to training, without using any data from the target recording. In the second strategy, validation was performed at the recording level. In each split, one recording was held out exclusively for testing, a second recording was reserved exclusively for validation, and the remaining eight recordings were used for training. All 90 ordered test–validation recording pairs were evaluated. The model was trained for up to 60 epochs. This strategy assessed the sensitivity of model selection to the choice of validation recording and provided an estimate of performance when validation and training data originated from different recordings.

#### 2.4.5 Baseline Models

To contextualize the performance of the proposed CNN–Transformer model, we compared it with both classical machinelearning and neural-network baselines. Classical methods were trained on handcrafted EMG features, including mean absolute value (MAV), root mean square (RMS), variance (VAR), and waveform length (WL), and included logistic regression, an RBF-kernel support vector machine (SVM), and gradient-boosted decision trees (XGBoost). Neural baselines comprised a simple 1D-CNN, a CNN-GRU model, and a Transformer-only architecture. All baseline models were evaluated using the same within-recording and leave-one-recording-out protocols as the proposed model.

#### 2.4.6 Ablation Study

To quantify the contribution of individual architectural components, we performed a series of controlled ablation experiments in which a single component of the proposed model was removed or modified while all remaining components and training settings were kept unchanged. The evaluated variants included removal of the Transformer encoder, trainable envelope extraction, positional encoding, and training-time data augmentation, as well as replacement of the proposed single-scale preprocessing block with the alternative multiscale variant.

#### 2.4.7 Evaluation Metrics and Statistical Analysis

Classification performance was evaluated using both accuracy and macro-averaged F1 score. Macro F1 was selected as the primary evaluation metric because it assigns equal weight to each digit class and is therefore sensitive to class-specific performance. For within-recording evaluation, metrics were first averaged over the five repeated splits for each recording and then summarized across recordings. For standard LORO evaluation, each held-out recording constituted one aggregation unit. For the held-out-validation analysis, results were first averaged over the nine validation-recording choices for each test recording and then summarized across test recordings. All analyses were implemented in Python using PyTorch, and identical preprocessing and evaluation pipelines were applied across all neural models unless stated otherwise. For the controlled benchmark and ablation analyses, paired differences were calculated at the recording level, with each held-out recording constituting one paired unit (*n* = 10). Mean paired differences and 95% percentile bootstrap confidence intervals were estimated using 10,000 resamples, and paired comparisons were assessed using the Wilcoxon signed-rank test. These analyses were considered exploratory because six recording-level units (M1– M6) originated from the same participant and were therefore not statistically independent participant observations. Consequently, the resulting intervals and p-values characterize variation among the available recordings and should not be interpreted as population-level participant inference.

## 3 Results

The developed CNN–Transformer model was first evaluated using the three experimental protocols described in Methods. We then compared the proposed architecture with classical and neural baseline methods and quantified the contribution of individual architectural components using controlled ablation experiments. Overall results are shown in Figures 3–6. Figure 3 depicts a comparison of macro F1 scores across all evaluated training and adaptation settings. Recording-specific macro F1 scores across representative experimental settings are shown in Figure 4; Figure 5 includes aggregated confusion matrices for the proposed model in within-recording and cross-recording settings; and Figure 6 shows the trade-off between average classification performance and inter-recording variability.

**Figure 3:**
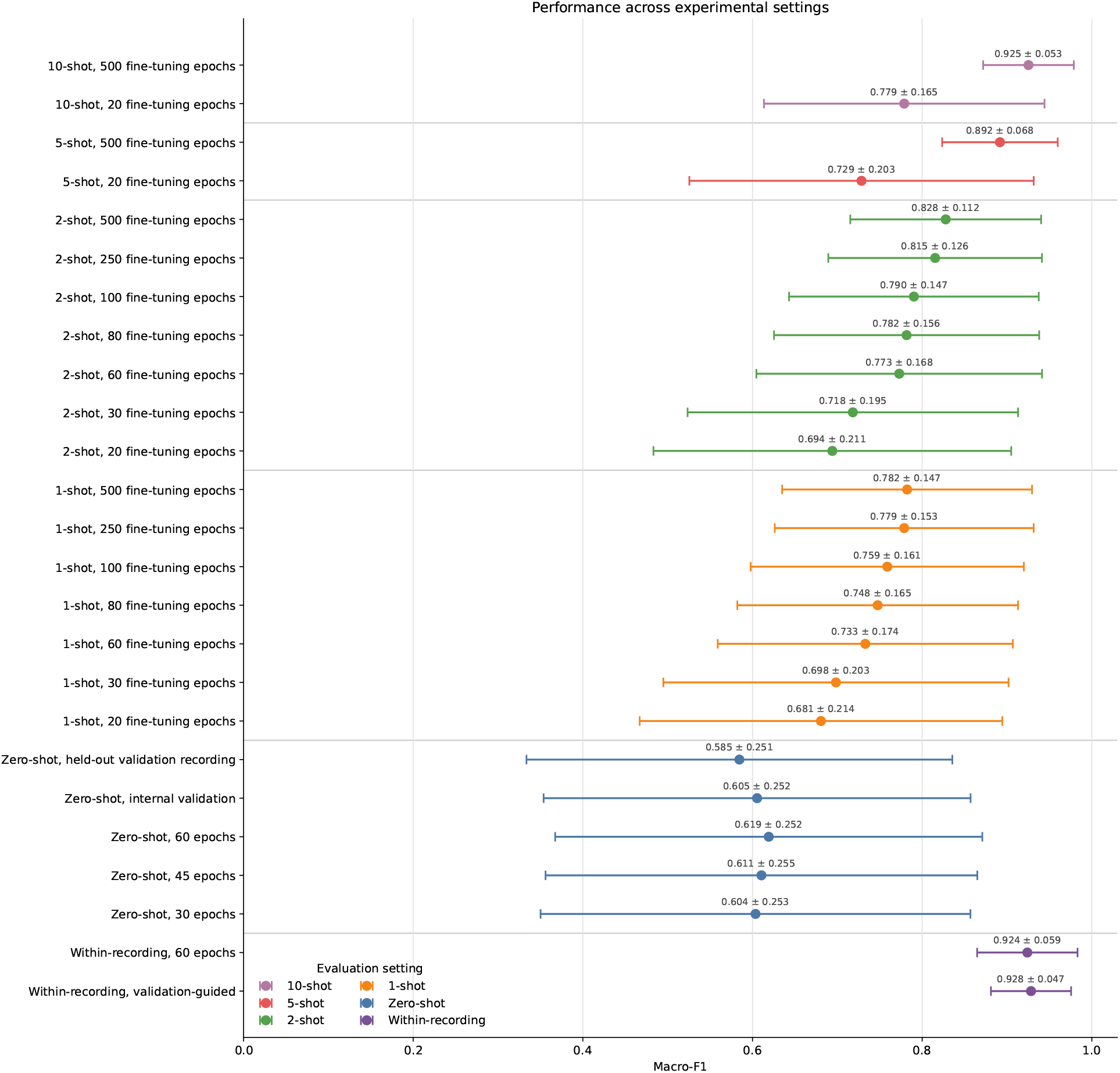
Comparison of macro F1 scores across all evaluated training and adaptation settings. Points indicate the mean macro F1 score across recordings, while horizontal error bars show the corresponding inter-recording standard deviation. The figure highlights the performance gap between within-recording and cross-recording evaluation and demonstrates the progressive improvement obtained through recording-specific fine-tuning.

**Figure 4:**
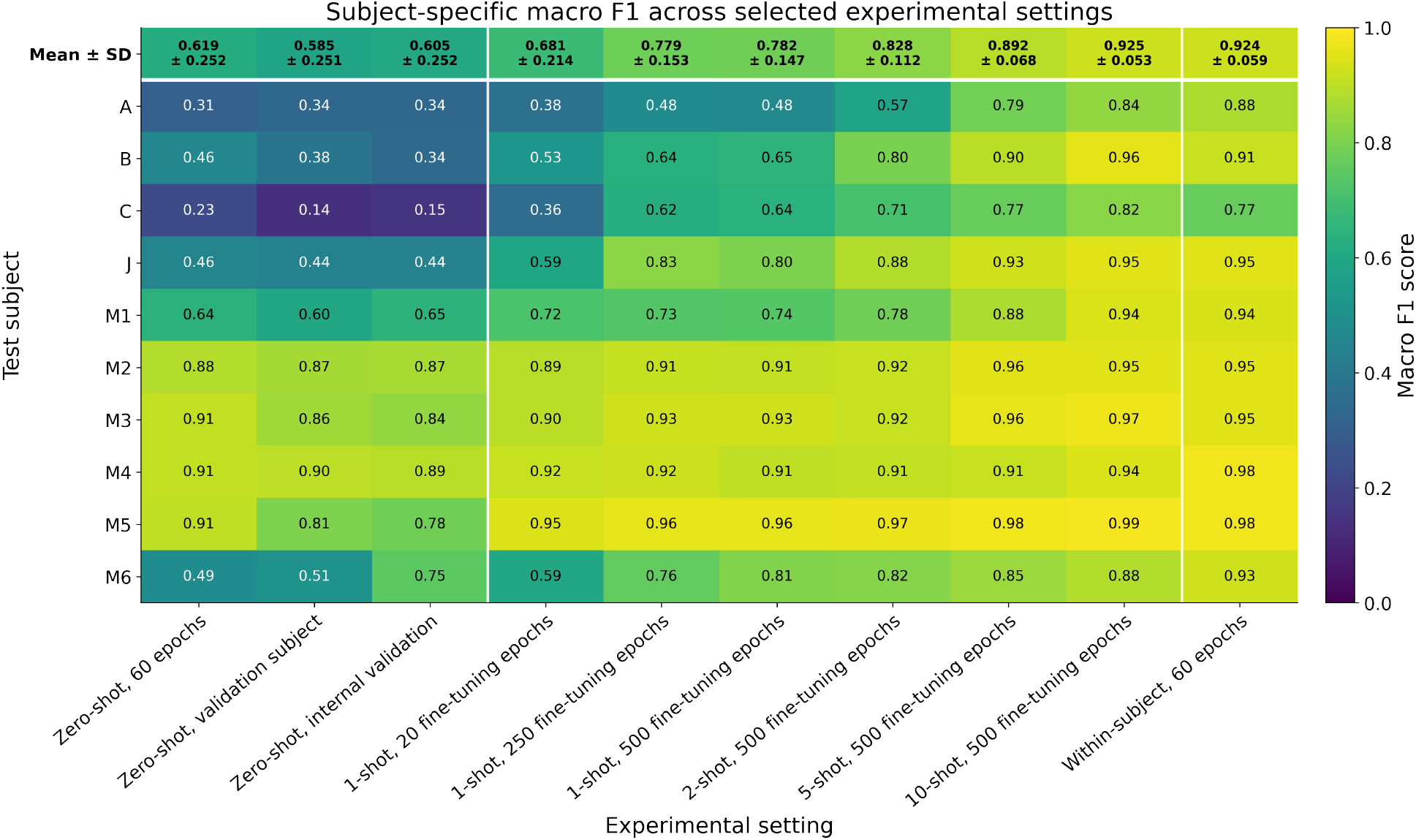
Recording-specific macro F1 scores across representative experimental settings. Each row corresponds to a test recording and each column to a training or adaptation strategy. Cell values report macro F1 scores. The heatmap reveals substantial interrecording variability and shows that recording-specific fine-tuning improves performance most strongly for recordings with poor zero-shot generalization.

**Figure 5:**
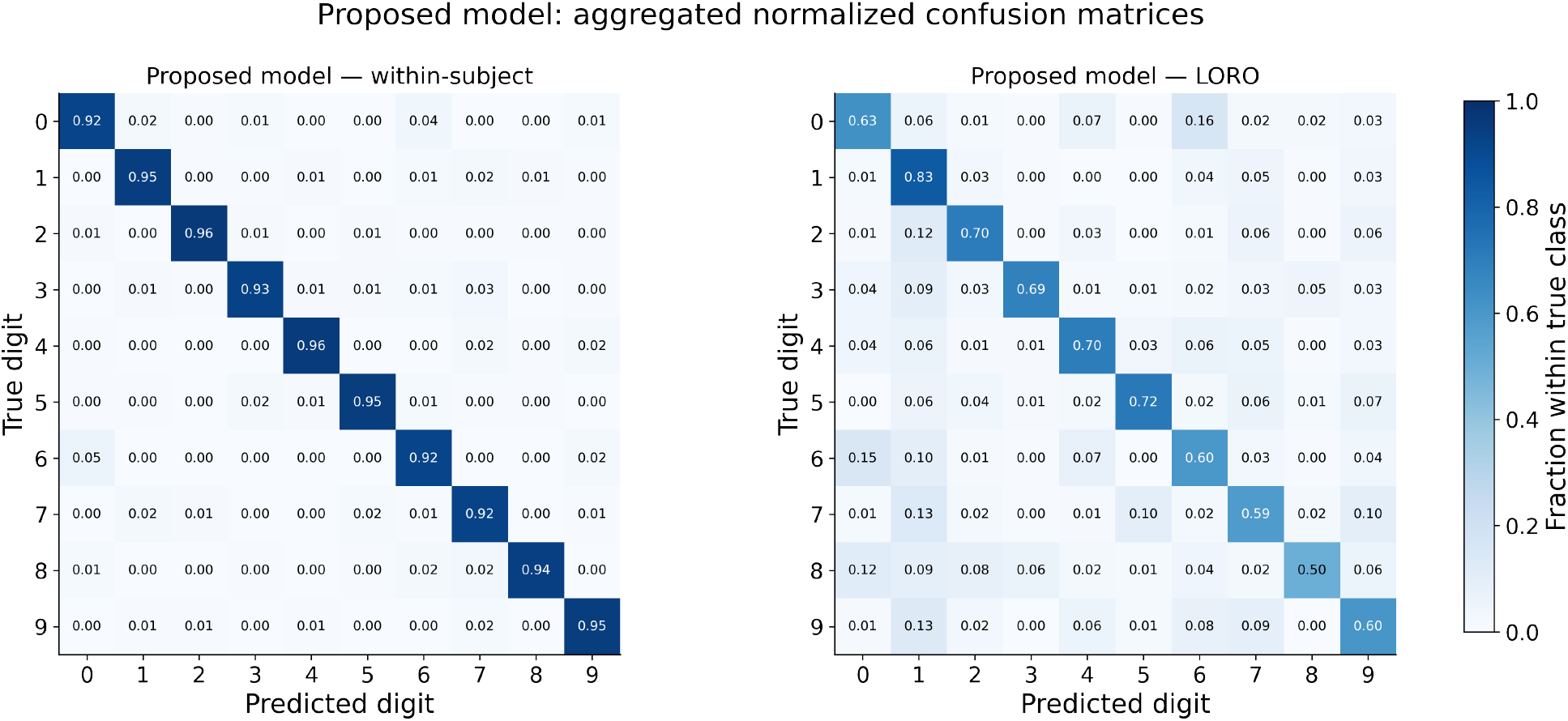
Aggregated confusion matrices for the proposed model. Row-normalized confusion matrices for the proposed single-scale CNN–Transformer under within-recording (left) and LORO cross-recording (right) evaluation. Matrices were aggregated across all recordings and folds before normalization. Each cell shows the proportion of trials from the corresponding true digit assigned to each predicted class.

**Figure 6:**
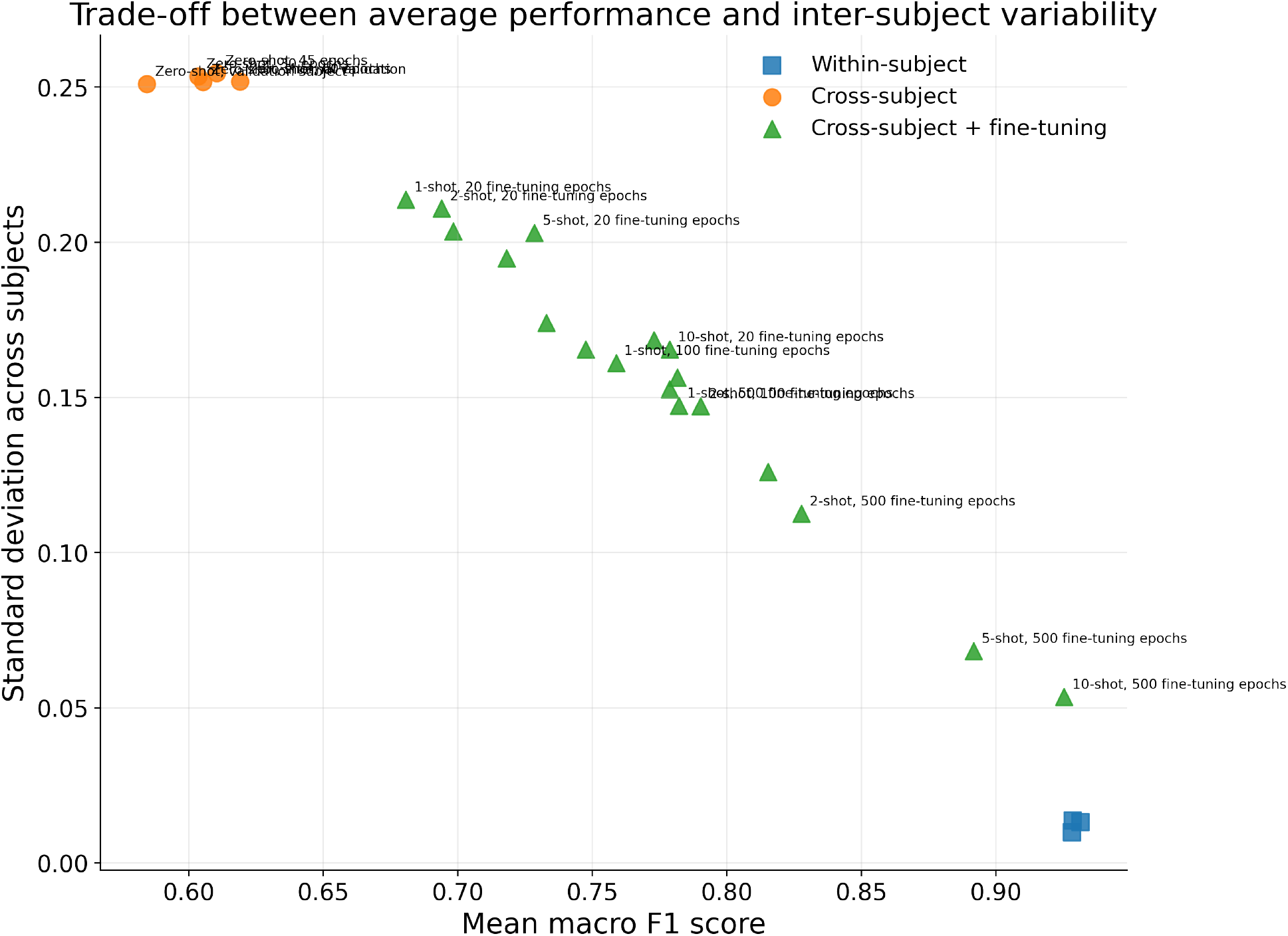
Trade-off between average classification performance and inter-recording variability. Each point represents one experimental configuration, with mean macro F1 on the horizontal axis and its standard deviation across recordings on the vertical axis. Desirable configurations are located toward the lower-right region, corresponding to high average performance and low inter-recording variability.

### 3.1 Within-Recording Evaluation

The within-recording evaluation quantified performance when all evaluated recordings were represented in the training data. With fixed 60-epoch training, the proposed single-scale CNN–Transformer achieved a mean macro F1 score of 0.924 *±* 0.059 across recordings, demonstrating that it achieved high classification performance when each evaluated recording was represented in the training data. In comparison, validation-guided model selection with early stopping achieved a slightly higher mean macro F1 score of 0.928 *±* 0.047, indicating a small numerical improvement in this setting.

### 3.2 Leave-One-Recording-Out Evaluation and Validation Strategies

The leave-one-recording-out (LORO) evaluation in the zero-shot setting achieved a mean macro F1 score of 0.619 *±* 0.252 without validation after 60 training epochs. The validation strategy influenced cross-recording performance. Selecting model checkpoints using validation trials sampled from the training recordings resulted in a mean macro F1 score of 0.605 *±* 0.252, whereas validation based on a separate held-out recording yielded 0.585 *±* 0.251. The validation-recording heatmap (Figure 7) showed considerable variability depending on the selected validation recording. Performance differences were particularly pronounced for recordings that already exhibited weak zero-shot generalization, indicating that a single validation recording may not adequately represent the variability of unseen recordings.

**Figure 7:**
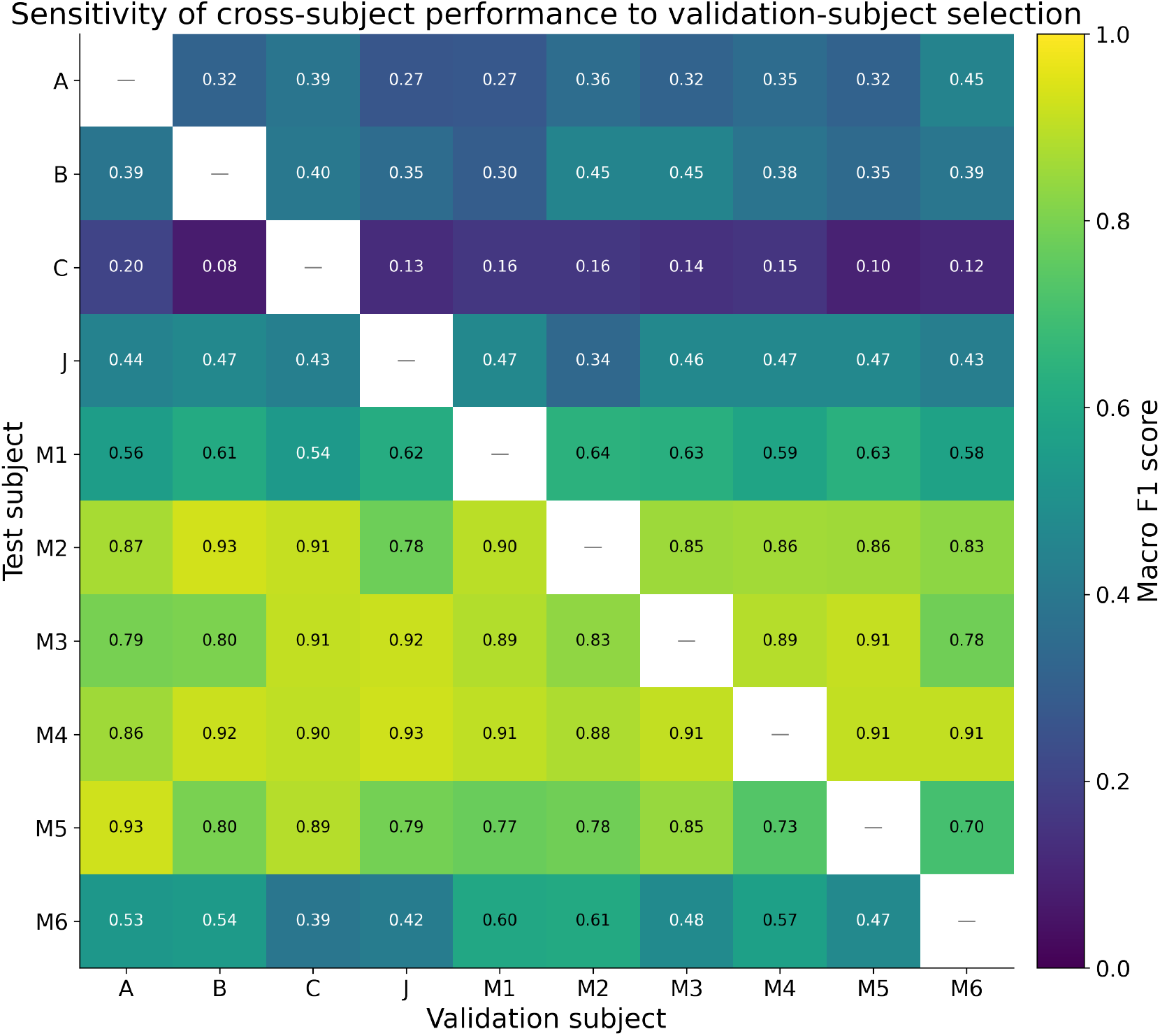
Sensitivity of cross-recording performance to the choice of validation recording. Rows denote test recordings and columns denote recordings used for validation. Cell values report macro F1 scores. The heatmap shows that model selection based on a single validation recording can substantially affect performance and that this sensitivity differs across test recordings.

**Figure 8:**
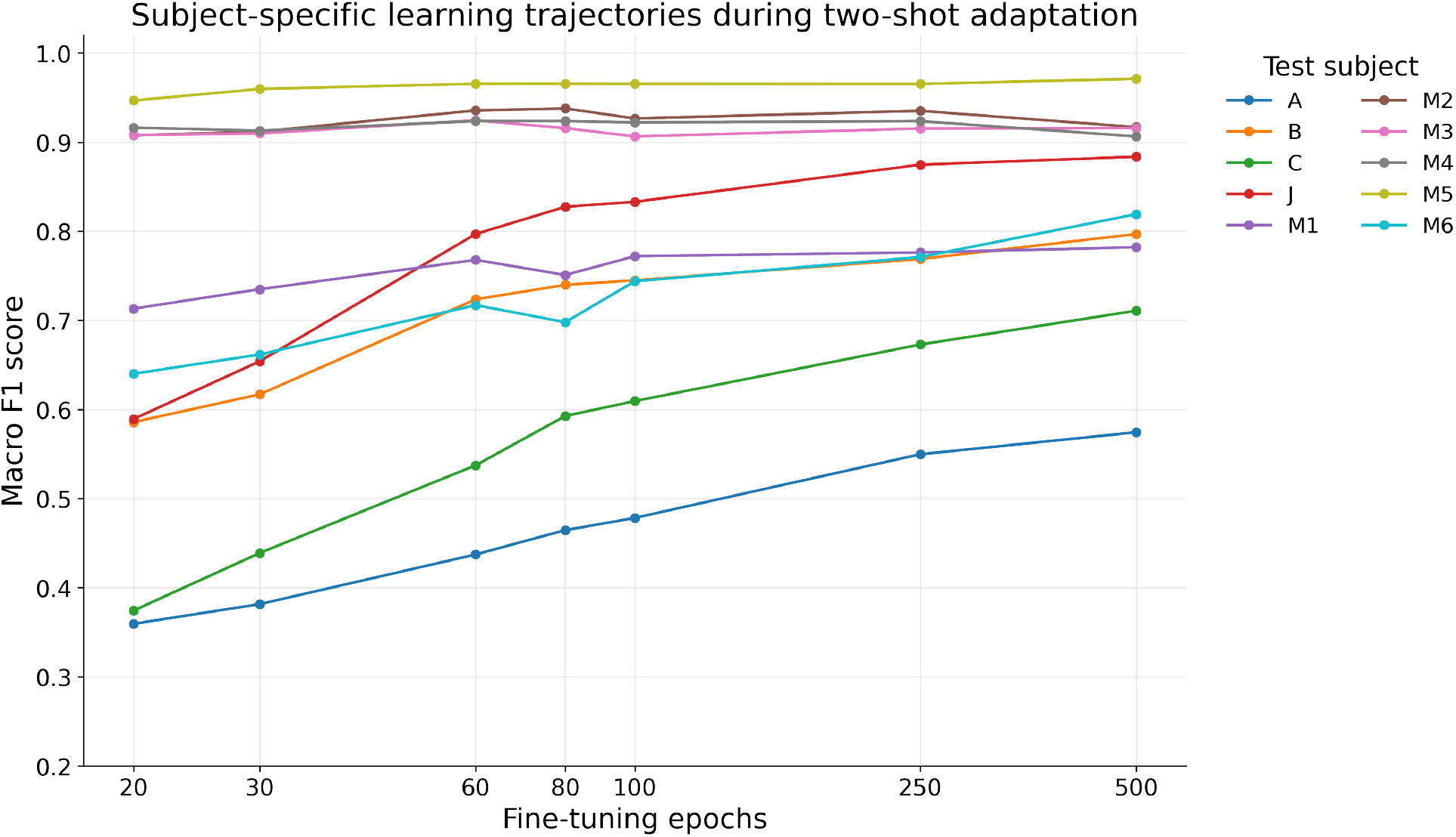
Recording-specific learning trajectories during two-shot fine-tuning. Each line represents the macro F1 score of one test recording across fine-tuning durations. The trajectories demonstrate that recordings benefit from adaptation at different rates, with difficult recordings showing larger and more gradual improvements than recordings that already achieve strong zero-shot performance.

**Figure 9:**
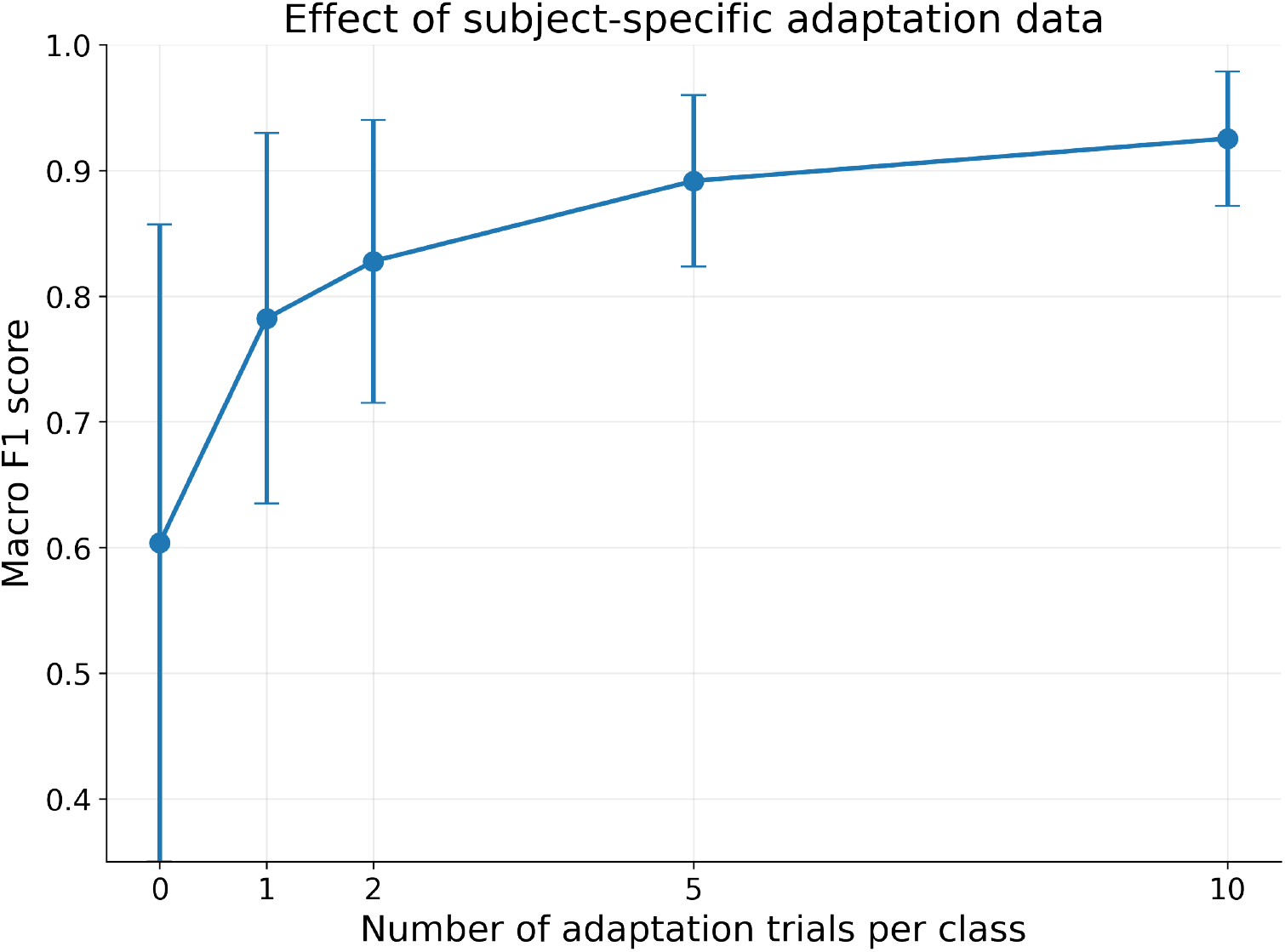
Effect of the number of recording-specific adaptation trials on cross-recording classification performance. Mean macro F1 is shown as a function of the number of adaptation trials per class. Error bars indicate the standard deviation across test recordings. Performance increases rapidly after introducing only a small amount of recording-specific data and continues to improve with additional adaptation examples.

### 3.3 Few-Shot Adaptation

Recording-specific fine-tuning consistently improved cross-recording performance. Even a minimal amount of labeled data from the target recording produced a noticeable improvement over the zero-shot baseline. With one adaptation trial per digit, macro F1 increased from 0.619 *±* 0.252 to 0.681 *±* 0.214 after 20 fine-tuning epochs and reached 0.782 *±* 0.147 after 500 epochs. Increasing the support-set size produced further gains. Two adaptation trials per digit increased the mean macro F1 score to 0.828 *±* 0.112, while five and ten trials per digit achieved 0.892 *±* 0.068 and 0.925 *±* 0.053, respectively. Thus, ten labeled examples per digit reached performance comparable to the within-recording setting. The recording-level heatmaps further demonstrated that adaptation was particularly beneficial for recordings with poor zero-shot performance. For example, recording B improved from 0.46 to 0.80 and recording J improved from 0.46 to 0.88 after only two-shot adaptation. Recordings with already high zero-shot performance, such as M2–M5, exhibited smaller absolute improvements because their initial performance was already close to the within-recording level.

### 3.4 Baseline Models

A separate benchmark compared the proposed CNN–Transformer with classical machine-learning and neural-network baselines (Figure 10). Within recordings, the proposed model achieved a macro F1 score of 0.935. 1D-CNN achieved 0.876 macro F1, and CNN-GRU achieved 0.811 macro F1. The Transformer-only model achieved a macro F1 score of 0.805. Performance among the classical baselines was heterogeneous: XGBoost achieved 0.839, whereas RBF-SVM and logistic regression achieved 0.679 and 0.503, respectively. The ranking changed substantially under the LORO protocol. The proposed CNN–Transformer achieved the highest cross-recording performance (0.630), outperforming all baseline methods. Among the neural baselines, CNN–GRU, Transformer-only, and the simple CNN achieved macro F1 scores of 0.261, 0.253, and 0.131, respectively. Classical methods generalized similarly poorly, with macro F1 scores ranging from 0.163 to 0.177. Overall, the benchmark indicates that although simpler models can model recording-specific EMG patterns, the proposed CNN–Transformer learns substantially more transferable representations.

**Figure 10:**
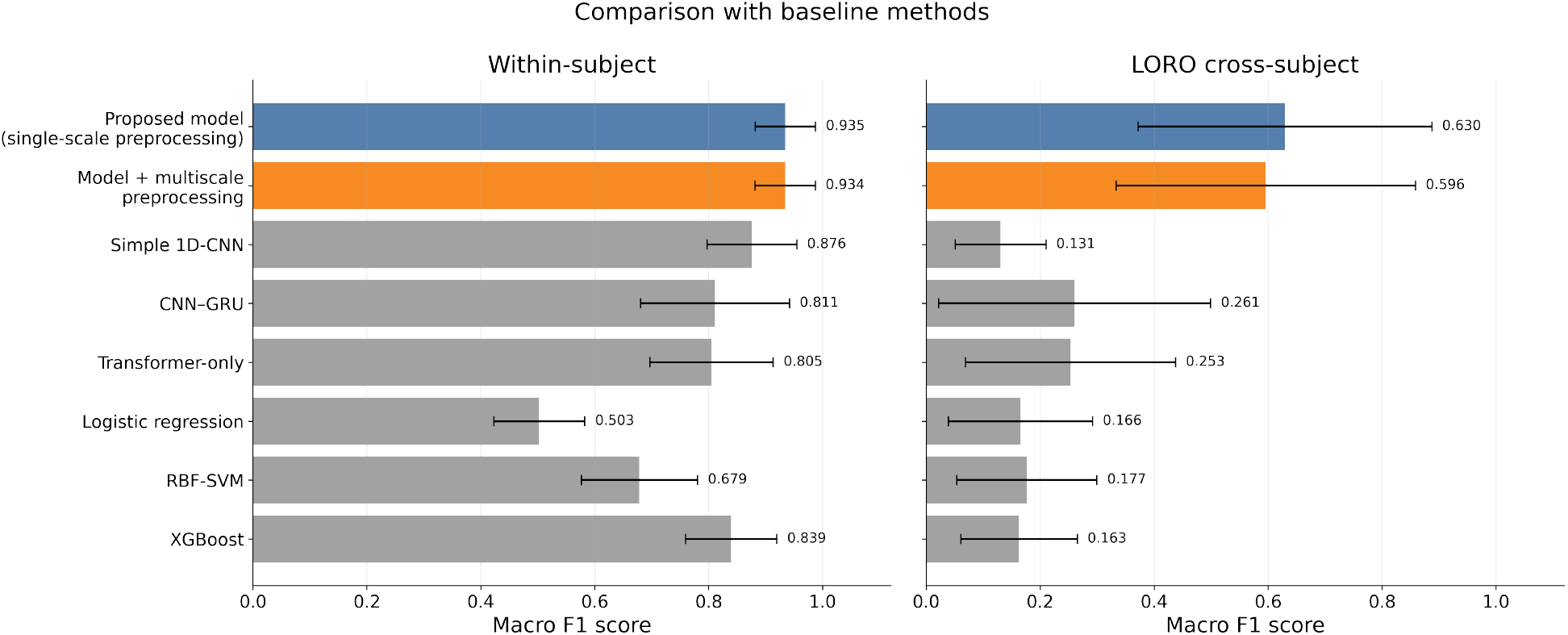
Comparison with baseline methods. Macro F1 performance of the proposed single-scale CNN–Transformer, its multiscale variant, and classical and neural baselines under within-recording (left) and LORO cross-recording (right) evaluation. Bars show mean macro F1 and error bars indicate standard deviation across recordings.

**Figure 11:**
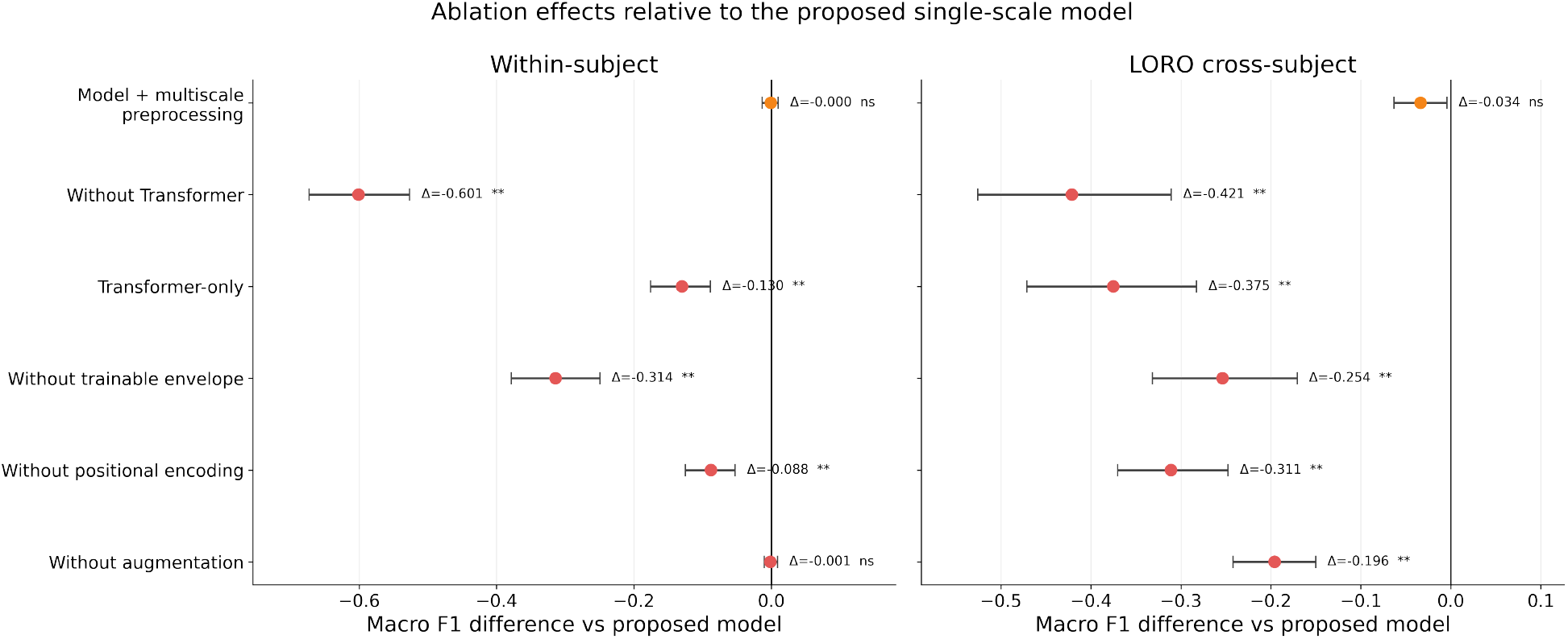
Ablation effects relative to the proposed model. Mean paired change in macro F1 after modifying or removing individual model components under within-recording (left) and LORO (right) evaluation. Points show mean recording-level differences, and error bars show 95% bootstrap confidence intervals. Positive values indicate improvement over the proposed model. Significance was assessed using the paired Wilcoxon signed-rank test: *\*p <* 0.05, *\*\*p <* 0.01; *ns*, not significant. The reported p-values were not adjusted for multiple comparisons and are presented for exploratory purposes.

**Figure 12:**
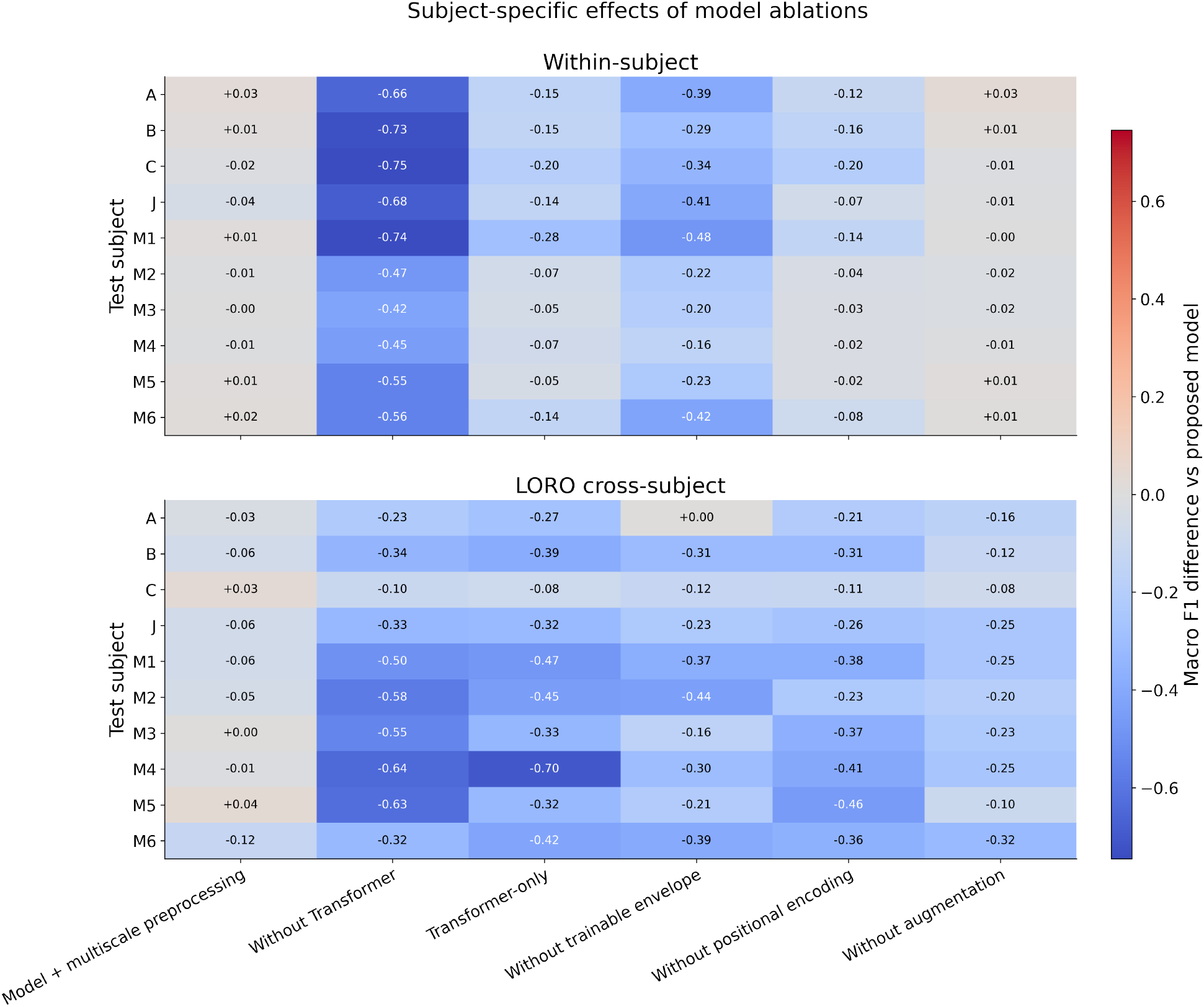
Recording-specific effects of model ablations. Change in macro F1 relative to the proposed single-scale model for each test recording under within-recording (top) and LORO cross-recording (bottom) evaluation. Red values indicate improved performance, whereas blue values indicate reduced performance.

**Figure 13:**
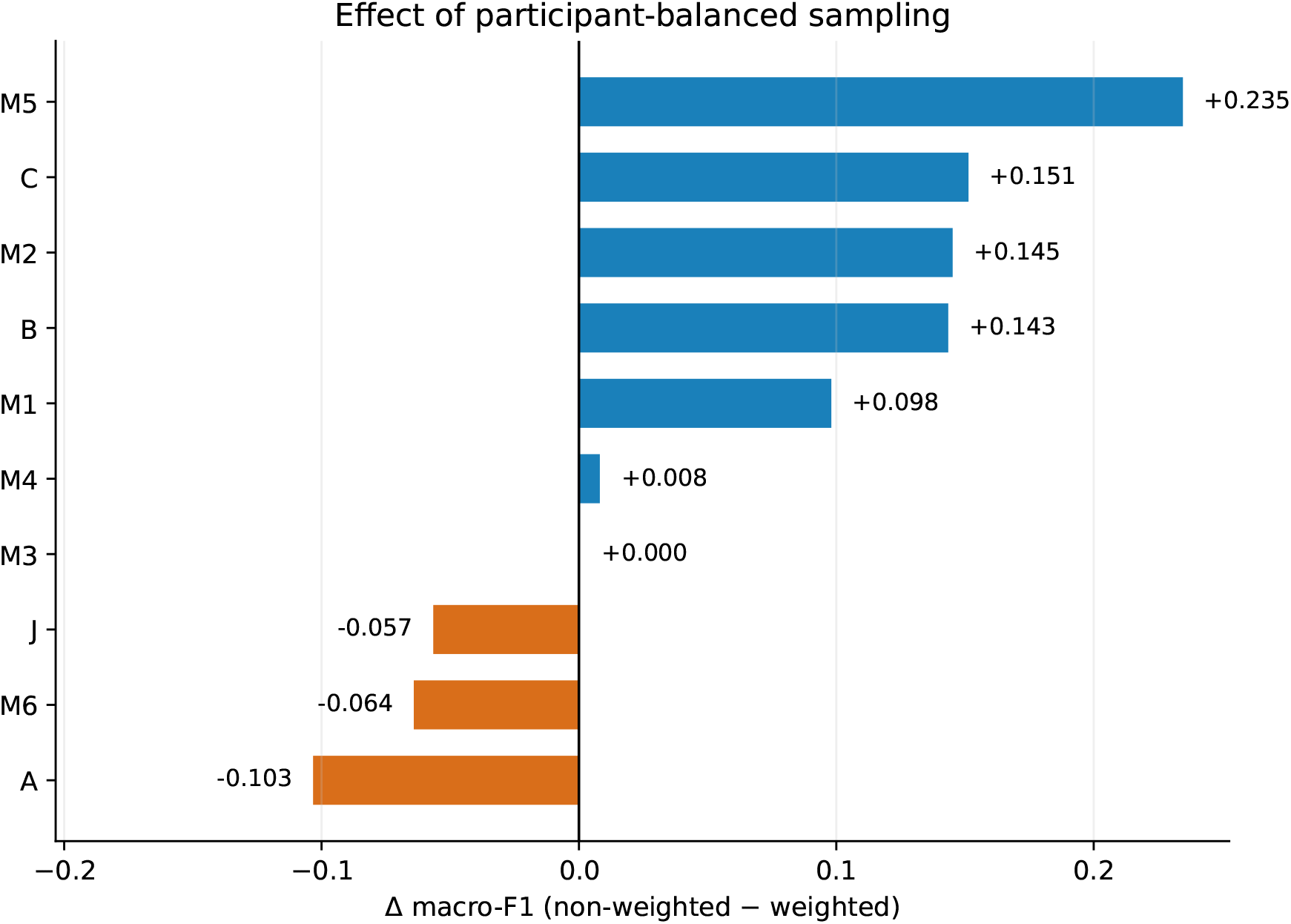
Effect of participant-balanced sampling on leave-one-recording-out (LORO) performance. Differences in macroaveraged F1 score between the non-weighted and participant-weighted training strategies are shown for each held-out recording (Δmacro-F1 = non-weighted - participant-weighted). Positive values indicate better performance with non-weighted sampling, whereas negative values indicate an improvement with participant-weighted sampling. Recordings A, B, C, and J correspond to independent participants, whereas M1–M6 represent repeated sessions from participant M. Participant weighting substantially reduced performance for several recordings, particularly M5, C, M2, and B, while improvements were observed for A, M6, and J; performance for M3 remained unchanged.

### 3.5 Ablation Study

The ablation study evaluated the contribution of the main architectural components relative to the proposed single-scale model. Removing the Transformer encoder caused the largest performance decrease: −0.601 macro F1 in the within-recording setting and −0.421 in the LORO setting. This confirms that long-range temporal modeling is especially important for cross-recording generalization. The Transformer-only variant also performed worse than the full model, with a decrease of −0.130 macro F1 within-recording and −0.375 in the LORO setting. This indicates that the Transformer encoder alone is insufficient and benefits from the convolutional front-end, which performs local temporal feature extraction and channel mixing before attention-based sequence modeling. Removing the trainable envelope extraction stage led to a decrease of −0.314 macro F1 within recordings and −0.254 in LORO evaluation. Removing positional encoding also reduced performance, with effects of −0.088 within recordings and 0.311 in LORO. These results suggest that both envelope-like preprocessing and explicit temporal position information contribute meaningfully to decoding performance. The effect of data augmentation differed between evaluation settings. In the within-recording benchmark, removing augmentation did not affect overall results (−0.001 macro F1), suggesting that augmentation was not necessary when training and testing data came from the same recordings. In contrast, in the LORO setting, removing augmentation reduced macro F1 by −0.196, indicating that augmentation was important for improving robustness to inter-recording variability.

#### 3.5.1 Participant-Weighted vs. Non-Weighted

To assess whether the six recordings from participant M disproportionately influenced model training, we performed a control analysis using participant-balanced sampling. During training, sampling weights were adjusted so that each of the five participants contributed equal total sampling probability, while the primary analysis sampled trials according to their original recording-level distribution. Under the LORO protocol, non-weighted sampling yielded higher macro F1 for six recordings (M5, C, M2, B, M1, and M4), whereas participant-weighted sampling improved performance for three recordings (A, M6, and J); performance for M3 remained unchanged. Because M1–M6 originated from the same participant and both strategies were evaluated using a single random seed, these results were interpreted as a descriptive control rather than a formal test of statistical significance.

#### 3.5.2 Single-Scale vs. Multiscale Preprocessing

The single-scale variant yielded numerically higher mean macro F1 scores in both benchmark protocols. In the baseline comparison, the single-scale model achieved 0.935 macro F1 in the within-recording setting and 0.630 in the LORO setting, whereas the multiscale model achieved 0.934 and 0.596, respectively.

The ablation analysis showed that replacing the single-scale preprocessing block with the multiscale block changed performance by −0.001 macro F1 in the within-recording setting and by −0.034 macro F1 in the LORO setting. Both effects were marked as not significant. Thus, the additional parallel convolutional branches did not provide a reliable improvement for the present dataset, and the simpler single-scale preprocessing block was used as the main model variant.

## 4 Discussion

This study evaluated a compact single-scale CNN–Transformer for decoding isolated handwritten digits directly from multichannel sEMG, with an emphasis on transfer to unseen recordings and low-data adaptation. Three main findings emerged. First, the model achieved high within-recording performance, reaching a mean macro F1 of 0.924 *±* 0.059. Second, zero-shot leave-one-recording-out (LORO) performance was substantially lower and more variable, with a mean macro F1 of 0.619 *±* 0.252. Third, a small amount of labeled data from the target recording substantially reduced this gap: two-shot adaptation reached 0.828 *±* 0.112, while ten-shot adaptation reached 0.925 *±* 0.053. In the separate controlled benchmark, the proposed model also provided the strongest LORO performance among the evaluated classical and neural baselines. Together, these results indicate that the primary model contribution is not only its ability to fit recording-specific EMG patterns, but also its comparatively strong transfer and efficient adaptation to previously unseen recordings.

The difference between within-recording and LORO performance illustrates the magnitude of the distribution shift encountered in practical sEMG systems. When training and test trials originated from the same recording, the model learned stable relationships between muscle activation patterns and digit classes. In contrast, a held-out recording may differ in signal amplitude, electrode contact, forearm posture, writing dynamics, muscular co-contraction, and temporal execution of individual strokes. Such variability is a recognized challenge in cross-user myoelectric pattern recognition [8]. The large LORO standard deviation is therefore an important result in itself: an average macro F1 of approximately 0.62 concealed a wide range of recording-level outcomes, indicating that zero-shot reliability depended strongly on the target recording.

The interpretation of cross-recording performance requires consideration of the dataset structure. Four participants contributed one recording each, whereas participant M contributed six recordings acquired on different days. The LORO protocol therefore combined two related but distinct transfer conditions: transfer between participants and transfer between sessions of the same participant. The comparatively strong zero-shot results for several M recordings suggest that day-to-day transfer within one participant may be less difficult than transfer to an entirely different participant. However, this distinction cannot be established conclusively because repeated sessions were available for only one participant. The participant-balanced sampling control further showed heterogeneous recording-level effects: non-weighted sampling performed better for six recordings (M5, C, M2, B, M1, and M4), participant-weighted sampling performed better for three recordings (A, M6, and J), and M3 remained unchanged. Because this control used a single random seed, it should be interpreted descriptively rather than as formal evidence that participant imbalance has no effect.

Few-shot adaptation produced the clearest practically relevant improvement. Only two labeled trials per digit, corresponding to 20 target-recording examples in total, increased mean macro F1 from the zero-shot level to 0.83. Increasing the support set to ten trials per digit raised performance to 0.925, essentially matching the within-recording result. Adaptation also reduced variability across recordings, suggesting that personalization improved both average recognition quality and consistency. These results favor a calibration-efficient system design rather than a strictly calibration-free one. A pretrained model could provide immediate approximate decoding and then be refined using a brief guided calibration procedure performed when the device is first used or after the electrode configuration changes.

The benefits of adaptation were not uniform. Recordings with weak zero-shot performance, including B and J, exhibited large improvements after fine-tuning, whereas several M recordings began at a comparatively high level and showed smaller absolute gains. The learning trajectories also differed across recordings: some reached a plateau after relatively few optimization steps, while difficult recordings continued to improve over longer fine-tuning intervals. A fixed calibration schedule may therefore be inefficient. An applied system could instead monitor support-set performance and stop adaptation once improvement becomes negligible. The post hoc trajectories indicate that a single fine-tuning duration was not optimal for every recording. Future work should investigate target-specific stopping based on a separate calibration-validation subset or cross-validation within the support set.

The validation experiments further demonstrated that model selection under cross-recording shift was not straightforward. Validation trials drawn from the non-target training recordings produced a mean macro F1 of 0.605 *±* 0.252, which was essentially comparable to fixed 60-epoch LORO training. Using an entirely separate recording for validation reduced performance to 0.585 *±* 0.251, and the validation-recording heatmap showed that the selected validation recording could substantially change performance on the target recording. A single validation recording was therefore not necessarily representative of an arbitrary future recording. In practical terms, complex source-domain model-selection procedures could provide limited benefit when no target data are available. Once even a small labeled target support set becomes available, direct adaptation appears to be more useful than attempting to identify a universally representative validation recording.

The baseline comparison clarifies the architectural advantage of the proposed model. In the controlled within-recording benchmark, several approaches could learn useful EMG representations, although their performance varied. Under LORO evaluation, however, all classical feature-based methods and the simpler neural architectures degraded substantially, whereas the proposed CNN–Transformer achieved the highest macro F1 of 0.630. Because the benchmark used matched settings across models, these values should be interpreted comparatively within that experiment rather than as replacements for the main protocol results. The benchmark nevertheless indicates that strong within-recording recognition does not guarantee transferability. The CNN–Transformer appears to provide a more suitable balance between local feature extraction and sequence-level modeling.

To examine the contribution of the selected architectural components, we conducted an ablation study. Removing the Transformer encoder caused the largest degradation, reducing macro F1 by 0.601 within recordings and by 0.421 under LORO. At the same time, the Transformer-only model also performed substantially worse than the complete architecture, particularly in the LORO condition. Thus, attention-based sequence modeling did not replace the convolutional front-end; the two components served complementary roles. The convolutional stages extracted local temporal patterns and inter-channel combinations, whereas the Transformer integrated information across the complete handwriting sequence. Trainable envelope extraction and positional encoding also contributed to performance. The particularly large LORO decrease after removing positional encoding suggests that preserving the order of motor events becomes more important when signal morphology varies across recordings.

Data augmentation primarily improved robustness rather than within-recording fitting. Removing augmentation had almost no effect on within-recording macro F1 but reduced LORO performance by 0.196. The simulated perturbations— including amplitude changes, temporal shifts, channel loss, and masking—therefore appear to expose the model to variations that are relevant to unseen recordings. By contrast, the multiscale preprocessing block did not provide a reliable advantage. Its performance was almost identical to the single-scale model within recordings and slightly lower under LORO, with neither paired difference reaching significance. One possible explanation is that fixed-length resampling, the subsequent convolutional extractor, and Transformer-based temporal modeling already provided sufficient sensitivity to temporal variation, limiting the additional value of parallel preprocessing kernels. Given its slightly lower complexity and comparable or better performance, the single-scale variant was retained as the proposed model.

From an applied-system perspective, the model has several favorable properties. It contains approximately 104,000 trainable parameters, and temporal downsampling reduces each 2500-sample trial to approximately 313 tokens before self-attention. This design limits the cost of Transformer processing and makes deployment in a compact adaptive interface plausible. The most realistic use case is a silent, biosignal-based input channel that complements rather than replaces established interaction modalities. For example, sEMG handwriting could be combined with gaze, speech, tablet input, or assistive control systems to provide an additional text-entry mechanism when visual tracking or conventional mechanical input is undesirable. Nevertheless, real-time latency, memory consumption, and energy use were not measured on target wearable hardware, and practical deployability should therefore be confirmed experimentally rather than inferred from parameter count alone.

Several limitations constrain the conclusions. Recording-level observations were not fully independent because six of the ten recordings were repeated sessions from the same participant. The recording-level statistical comparisons should therefore be interpreted as exploratory rather than as confirmatory participant-level inference. The present LORO results consequently cannot fully separate cross-session from cross-participant generalization. The task was limited to isolated digits with offline trial boundaries derived from tablet signals; a real text-entry interface would require online detection of writing onset and offset, continuous sequence recognition, and mechanisms for correcting errors. All segments were resampled to a fixed length, and the robustness of the model to naturally variable-duration input remains to be evaluated. The experiments also did not systematically test repeated sensor donning, unconstrained arm posture, or long-term use. In addition, few-shot support sets were labeled and balanced across digits, which represents a more controlled calibration procedure than may be available in everyday operation.

Future work should therefore use larger participant-balanced datasets containing repeated sessions for every participant, allowing cross-session and cross-participant transfer to be evaluated separately. Self-supervised pretraining, domain-invariant representation learning, and explicit domain adaptation may further improve zero-shot generalization. Adaptive stopping criteria and active selection of calibration examples could reduce the time required for personalization. Finally, integration with online segmentation and evaluation on wearable hardware is necessary to determine whether the proposed approach can support continuous, real-time interaction.

Overall, the results show that a compact CNN–Transformer can decode handwriting-related sEMG with high within-recording performance and provides stronger cross-recording transfer than the tested alternatives. Zero-shot decoding remains sensitive to recording-specific variability, but few-shot adaptation recovers most of the lost performance using a limited number of labeled examples. This combination of transferable representation learning and lightweight personalization provides a practical foundation for future adaptive sEMG-based input systems.

## Acknowledgements

This work was supported by the The Ministry of Economic Development of the Russian Federation in accordance with the subsidy agreement (agreement identifier 000000C313925P4H0002; grant No 139-15-2025-012).

## References

[1] Jhansi Rani Gundala, Mohammad Farukh Hashmi, and Aditya Gupta. Surface electromyography and artificial intelligence for human activity recognition—a systematic review on methods, emerging trends applications, challenges, and future implementation. IEEE Access, 11:105140–105169, 2023.

[2] Jose Guadalupe Beltran-Hernandez, Jose Ruiz-Pinales, Pedro Lopez-Rodriguez, Jose Luis Lopez-Ramirez, and Juan Gabriel Avina-Cervantes. Multi-stroke handwriting character recognition based on sEMG using convolutional-recurrent neural networks. Mathematical Biosciences and Engineering, 17(5):5432–5448, 2020.

[3] Jose Ruiz-Pinales, Jose Guadalupe Beltran-Hernandez, Juan Gabriel Avina-Cervantes, Pedro Lopez-Rodriguez, and Jose Luis Lopez-Ramirez. HCMYO-A dataset, 2020. Dataset.

[4] Ashish Vaswani, Noam Shazeer, Niki Parmar, Jakob Uszkoreit, Llion Jones, Aidan N. Gomez, Łukasz Kaiser, and Illia Polosukhin. Attention is all you need. In Advances in Neural Information Processing Systems, volume 30, pages 5998–6008. Curran Associates, Inc., 2017.

[5] Yanhong Liu, Xingyu Li, Lei Yang, and Hongnian Yu. A transformer-based gesture prediction model via sEMG sensor for human–robot interaction. IEEE Transactions on Instrumentation and Measurement, 73:1–15, 2024. Art.no. 2510615.

[6] Mansooreh Montazerin, Soheil Zabihi, Elahe Rahimian, Arash Mohammadi, and Farnoosh Naderkhani. ViT-HGR: Vision transformer-based hand gesture recognition from high-density surface EMG signals, 2022.

[7] Stephen Butterworth. On the theory of filter amplifiers. Experimental Wireless & the Wireless Engineer, 7:536–541, October 1930.

[8] Xuan Zhang, L. Wu, Xu Zhang, Xiang Chen, Chang Li, and Xun Chen. Multi-source domain generalization and adaptation toward cross-subject myoelectric pattern recognition. Journal of Neural Engineering, 20(1):016050, 2023.

[9] Jake Snell, Kevin Swersky, and Richard S. Zemel. Prototypical networks for few-shot learning. In Advances in Neural Information Processing Systems, volume 30, pages 4077–4087. Curran Associates, Inc., 2017.

[10] Michael Linderman, Mikhail A. Lebedev, and Joseph S. Erlichman. Recognition of handwriting from electromyography. PLOS ONE, 4(8):e6791, August 2009.

[11] Ilya Loshchilov and Frank Hutter. Decoupled weight decay regularization.In International Conference on Learning Representations, 2019.

